# Analysing Patterns in Population Dynamics using Repeated Population Surveys with Three Types of Detection Data

**DOI:** 10.1101/459537

**Authors:** Guillaume Péron, Mathieu Garel

## Abstract

1. We generalize the distance sampling protocol to accommodate three types of detection data in population surveys (population counts): distance to the observer, multiple observers, and time-to-detection. We also account for the effect of a partially-observed individual covariate, with the aim to account for the non-independence of individuals in groups and the effect of group size on detection probability. Finally, we separate the probability of availability to detection and the probability of detection when available.
2. We compute the statistical power of our new model using simulations and illustrate some of the biases in the simpler alternative analytical procedures that oue new formulation allows to avoid. We discuss issues of weak identifiability (an issue shared with other state-space models) and the bias-precision tradeoff in population survey analyses.
3. We recommend maintaining both simple analyses of population survey data and more complex analyses of the detection process, maybe in a dashboard of indicators. Discrepancies between results from simple and complex analyses can help identify sources of biases and loss of precision.

## INTRODUCTION

Observed patterns of variation in animal abundance may yield a flawed picture of the underlying population dynamics because of imperfect sampling (Link & Sauer, 1998; Royle, 2004;Engeman, 2005). Typically, variation in observed abundance may be caused by variation in the probability to detect individual animals in the population. A broad range of methods have been proposed and successfully used to overcome this issue (Williams, Nichols, & Conroy, 2002). Our first objective here is to quickly review these methods, their domain of application, and some of their shortcomings when applied to more challenging species. Second, we aim to introduce a new framework designed to adress those shortcomings, mostly by adding a few generalizing features to the well-known distance sampling framework (S. Buckland, Anderson, Burnham, & Laake, 1993). Third, we use simulations and real case studies to thoroughly discuss the bias-precision trade-off and the issue of weak identifiability of model parameters. The bias-precision trade-off refers to the fact that adding more parameters to deal with sources of bias unavoidably leads to less precise estimates (Burnham & Anderson, 2002), calling for a careful weighing of the pro and cons of complexifying the analytical framework. Weak identifiability refers to the difficulty to separate the relative contributions to the variance in the data of different model parameters, leading to estimation issues when the data are sparse (Auger-Méthé et al., 2016; Barker, Schofield, Link, & Sauer, 2017).

## A QUICK REVIEW OF THE METHODS TO ANALYZE PATTERNS IN POPULATION DYNAMICS USING COUNT DATA

### The Index of population size methodology (IPS)

Hereafter the acronym “IPS” refers to methodologies that infer patterns in population dynamics using the expected count, i.e., the product between the population abundance and the probability of detection. Some IPS methods consist in averaging the count over several replicates, i.e., they “average out” the sampling variance around the expected count (Loison, Appolinaire, Jullien, & Dubray, 2006). These methods assume that the expected detection probability is the same everywhere and every time, and that most of the noise around the expected count is caused by counting errors.

Alternatively, one may rely on linear models of the count across space and time. Linear predictors and random effects would then control for factors of variation in detection probability, such as observer proficiency, vegetation type, or weather (Link & Sauer, 1998), thereby relaxing the assumption that the expected detection probability is the same everywhere and every time.

The main issue with the otherwise attractively simple and effective IPS methodologies is that, if a factor jointly influences population abundance and detection probability (Anderson, 2003), it will not be possible to tease apart these two influences. Furthermore, the factors of variation in detection probability may not be *a priori* known and quantified, preventing their inclusion as explanatory variable. Lastly, count data are often very noisy, in which case IPS methods can become unreliable or request too many replicates to be tractable (Harris, 1986; Gerrodette, 1987).

### Population reconstruction from individual-based data

Based on the above shortcomings of the IPS methodologies, researchers have increasingly preferred to “reconstruct” the population dynamics from estimates of vital rates, such as survival and fecundity (Caswell, 2001; Besbeas, Freeman, Morgan, & Catchpole, 2002; Williams et al., 2002). In this approach, one uses individual-based data to compute, each year, the balance between the births and deaths, and thereby the population growth rate, yielding an index of population abundance relative to the abundance at the start of the study. The main advantage of this approach is the ability to investigate individual and environmental variation in vital rates, and thereby obtain realistic models of population dynamics likely to yield reliable short-term predictions (Gauthier, Péron, Lebreton, Grenier, & Oudenhove, 2016). The main issue is the cost and field-intensiveness.

### Unmarked methods

To avoid the shortcomings of the IPS and population-reconstruction methodologies, the “unmarked” philosophy (Fiske & Chandler, 2011) is currently gaining in popularity. The “unmarked” philosophy refers to methods that do not require individual-based data from marked or otherwise recognizable individuals, but that still separate the variance in the count data into a sampling (detection) and a process (population dynamics) components. The most widely used “unmarked” methodology is distance sampling (S. Buckland et al., 1993). In distance sampling, the decline in recorded abundance with distance to the observer is leveraged to estimate the detection probability as a function of distance and correct the count for imperfect detection. Another seminal model underlying the “unmarked” philosophy is the N-mixture model (Royle,2004). The sampling variance is estimated by fitting a binomial model to the variation across replicates. These two classes of methods have been widely and successfully applied to a range of monitoring problems, but their widespread use should not hide that both methods are based on strong assumptions, which, when violated, lead to serious shortcomings.

In particular, the N-mixture approach is empirically well-known to yield overestimated or infinite estimates of population size when detection probability is small or there are few replicates (Couturier, Cheylan, Bertolero, Astruc, & Besnard, 2013; Dennis, Morgan, & Ridout, 2015). Recently, Barker et al. (2017) explained that the issue is linked to the difficulty to distinguish between the Poisson and binomial distributions of the variance across replicates when the data are sparse, whereas the assumption that the variance is binomial is critical to identify the parameters in the N-mixture model. More generally, state-space models, of which N-mixture models are a particular case, are known for their identifiability issues, i.e., when the sampling (detection) variance largely exceeds the process (population dynamics) variance, only the sum of the process and sampling variance is estimable (as nicely illustrated in Appendix E of Auger-Méthé et al., 2016). This issue is obviously of major concern since the unique goal of state-space modeling in this case is to separate the two variance components to get at the population dynamics. In addition, the N-mixture model assumes that the detection probability is constant across replicates, which arguably prevents the accurate description of the sampling variance (Barker et al., 2017). Lastly, the N-mixture model fitting procedure in the Bayesian framework is sensitive to the arbitrary choice of a maximum potential population size, but this issue can arguably be dealt with using *a priori* biological insights (Couturier et al., 2013; Dennis et al., 2015).

Now regarding the distance sampling methodology, the main constraint is that, at least in its initial formulation, the framework relied on the assumption that animals (or clusters of animals) are uniformly distributed within the field of vision of the observers (S. Buckland et al., 1993). This means that animals are equally likely to be present at any point in space. Although that assumption has since been partly relaxed (Conn, Laake, V Johnson, 2012; Sillett, Chandler, Royle, Kery, & Morrison, 2012; Hostetler & Chandler, 2015), the complex modeling and data density required to relax that assumption mean that most users still eventually work under that assumption that animals are uniformly distributed across the field of vision. Another issue with distance sampling is that hiding behaviors, vertical movements (diving, climbing trees), and more generally temporary emigration out of the survey area leads some individuals to be temporally unavailable to detection. They are still part of the population, but their detection probability is temporarily zero. Buckland et al. 1993 introduced the familiar *g_0_* term, a.k.a. “availability probability” to deal with this. But, *g_0_* and its variation in time and space must be documented separately, for example with telemetry data (e.g., Couturier et al. 2013). This requirement for external data can suppress the cost-effectiveness of the method. Another lingering criticism of distance sampling is that most of the readily available implementations are geared towards obtaining snapshots of the population abundance, not monitoring fluctuations in abundance over multiple years or sites. In particular, these implementations do not facilitate the borrowing of information across years and sites.

Circumventing all these issues, while remaining cost-effective and realistic in terms of data requirements is currently an area of active research (Chandler, Royle, & King, 2011; Conn et al., 2012; Sillett et al., 2012; Dénes, Silveira, & Beissinger, 2015; Hostetler & Chandler, 2015; Clement, Converse, & Royle, 2017). The key is to collect auxiliary data during the counts. We call these auxiliary data the “detection data”.

## OUR MODEL

The model was inspired by surveys of mountain ungulate populations in France (gregarious mammals that live in rough terrain with impaired observer visibility), but we expect it to be relevant in other situations as well. We first review the three types of “detection data” that we consider, then we describe the likelihood function that allows their joint analysis, and finally we describe a few necessary post-hoc manipulations of parameter estimates.

### Three types of detection data for unmarked animals

The first type of detection data is distance to the observer, i.e., our model is a generalization of the distance sampling protocol.

The second type of detection data comes from the multiple-observer protocol (Nichols, Hines, Sauer, V Fallon, 2000). Several observers perform the same counting procedure. For each detected individual, the series of detection or non-detection by each observer generates an history of detection akin to a capture-recapture history (but with the distance to the observer recorded as covariates). In a nutshell, the proportion of observers that detected an individual informs the detection probability of that individual. Some logistical issues however prevent the implementation of a full multiple-observer protocol. Observers indeed typically influence each other: when an observer records an animal, other observers are likely to notice and subsequently detect the same animal using cues unwillingly provided by the first observer. For this reason, we advocate (and we implement in our model) a removal design for the multiple-observer protocol. The removal design means that we establish an order among the observers. Observer n+1 can only add new detections that observer *n* did not make. In addition to avoiding the above-described positive bias in the detection probability of observer n+1, the removal design requires less communication between observers than the full multiple-observer protocol and is thus more straightforward to implement.

The third and last type of detection data is generated by a time-to-detection protocol (Alldredge et al. 2007; a.k.a. removal sampling protocol *sensu* Fiske and Chandler 2011). For this protocol, we assume that the time to detection scales to the instant detection probability. In practice, we may discretize the detection process by dividing the count period into secondary occasions. Then, the series of detections and non-detections at each secondary occasion constitutes a capture-recapture history for each detected individual. However, once an individual has been detected once, its probability of detection is drastically improved because the observers now know that this individual is present and roughly where it is. For this reason, we also implement a removal design for the time-to-detection protocol. We only record the first secondary occasion at which an individual is observed.

The joint analysis of different types of detection data has already been proposed previously. For example, Chandler et al. (2011) described how to combine the multiple-observer and time-to-detection protocols to estimate abundance in the presence of temporary emigration. Conn et al. (2012) described how to combine the multiple-observer and distance sampling protocols to estimate spatial variation in population density. Here we combine the three types of detection data, in a flexible manner, meaning that, in some sites or years, users might decide to apply only one, two or the three protocols, and still be able to analyze the resulting data within the same framework. To represent temporary emigration out of valleys or mountain slopes by our focal mountain ungulates, we consider temporary emigration like Chandler et al., but unlike Conn et al. we still use discrete geographic units (sites) to represent spatial variation in abundance, in part because in mountain ungulate monitoring schemes, sites are relatively small (walkable in a few hours) and homogeneous (grazing grounds).

### Model likelihood

Because mountain ungulates (our motivation for the new development) often live in groups, the statistical unit in our model is the group of animals, or the cluster *sensu* Buckland et al. (1993). One of our concerns is the effect of group size on detection probability, and in particular the way in which covariation between abundance and group size may flaw the IPS methodology. In other words, if group size increases with abundance (Toïgo, Gaillard, & Michallet, 1996; Pépin & Gerard, 2008), and detection probability increases with group size, the observed population growth rate may be artificially inflated, potentially leading to over-optimistic management decisions. We consider that, during any visit, the groups are either available or unavailable. Each detected group is described by two group covariates: the group size and the distance to the observer. The distance may be recorded exactly, or binned into classes of approximate distance. The group size is considered error-free; there is no counting error or partial availability of groups. To deal with counting errors or partial availability of groups, see Clement et al. (2017), but this feature is not supported in our framework.

As described under “Three types of detection data” above, we use a removal design for both the multiple-observer and the time-to-detection protocols. This means that we record the first time an individual was detected (and not whether it was detected again afterwards), and the first observer in an ordered series of observers that recorded the individuals (and not whether the observers that came afterwards in the series of observers detected it). We acknowledge that implementing a removal design instead of a full capture-recapture design induces some loss of information, but as argued under “Three types of detection data”, this information would likely be flawed.

We denote θ the set of model parameters (Table 1) and Ythe detection data. *Y* is stratified across *K* sites, *T*years, *U_k,t_* within-year visits to site *k*in year *t, V_k t,u_* secondary occasions within visit *u* to site *k*in year t, and *O_k,t,u_*observers. As noted above, *U_k,t_, V_k,t,U_*, and *Ok.t.u* can change across sites, years, and visits, allowing for a flexible study design. For example *Ok,t,u ^=^ 1* means that only one observer participated in the survey of site k, year t, and visit u.The likelihood *L(θY)* describes the probability to record *Y* as a function of θ. For each detected group i, we know the site k, the year t, the number of the visit u, the secondary session, the observer *o_t_*, the distance d_i_, and the group size *g_i_.* From these data we can compute the probability P_*k,t,U,i*_ that the group was detected, as the product of four terms: the probability that the group was available for detection, the probability that it was not detected until observer *o_i_* the probability that observer *o_t_* did not detect it until subsession *v_i_*, and eventually the probability that the observer detected it.

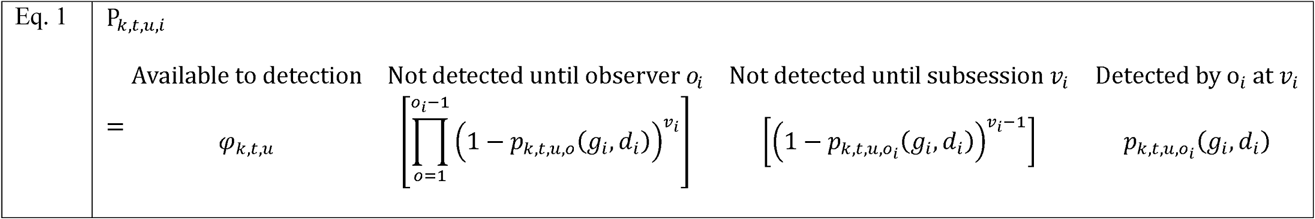

**Table 1:**
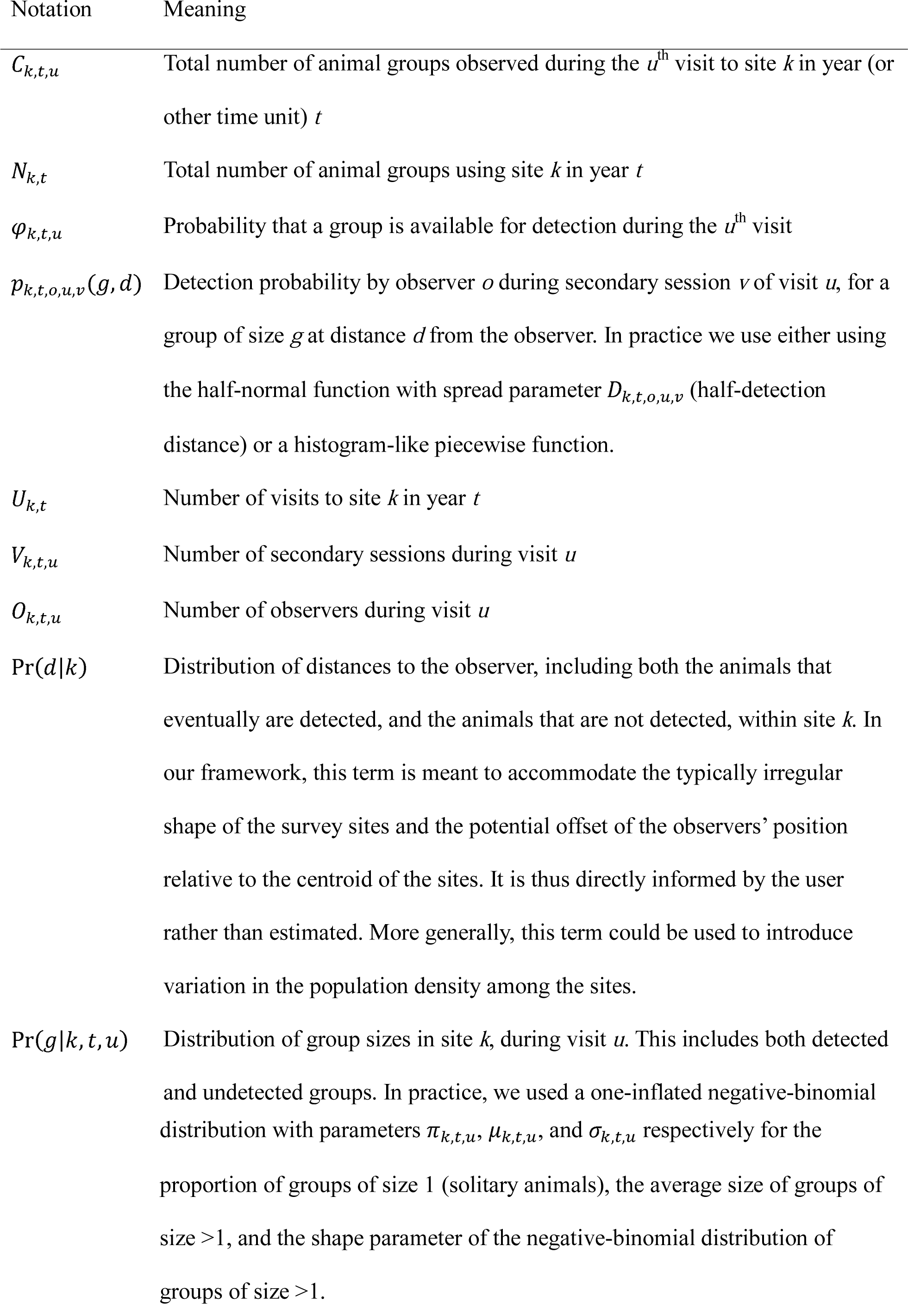
Notation for the ‘chamois’ class of models

The product between the first pair of brackets is replaced by a one if *o_i_* = 1. All notation is summarized in Table 1. The product of all the *P_k.t.u.i_* terms corresponds to the overall probability to detect the groups that were detected, in the way they were detected.

Then we need to account for the groups that were not detected. This is where the population abundance actually enters the likelihood, because the population abundance in site k and year t is the sum of the number of individuals recorded during any visit to site k in year t, plus the number of individuals that escaped detection during that visit. The challenge is that the group size and distance from the observer are, obviously, not known for individuals that were not detected. We tackled this as a simple extrapolation problem. We introduced the distribution of distance to the observer, denoted Pr(d|k), and the distribution of group sizes, denoted *Pr(g|k, t, u).* We assumed that both detected and non-detected groups were drawn for the same distributions. For Pr(d|c), we further assumed that the groups were uniformly distributed within each site, so that Pr(d|c) only depended on the configuration of the site, as informed by a field of view analysis in a GIS software. For *Pr(g|k, t,u)*, we used a one-inflated negative-binomial distribution of group sizes. The three parameters of that distribution (Table 1) were part of the list of parameters to be estimated, θ. We chose the one-inflated negative-binomial distribution rather than alternatives (e.g., Poisson) based on recommendations by Ver Hoef & Boveng, (2007) and on the observation that there was an excess of solitary animals relative to the negative-binomial distribution in real data. It is theoretically possible to replace this distribution with a double-Poisson distribution (Barker et al., 2017) but that is not implemented. Lastly, we assumed, as is commonly verified in distance sampling applications (Miller, 2015), that the link between detection probability and distance followed a half-normal function. The spread parameter a.k.a. half-detection distance of that function, denoted *D_k t o u_*, was made to vary log-linearly with group size. The result was the function *P_k,t,u,o_(9, d)* giving the site-, year-, visit-, and observer-specific detection probability as a function of group size and distance to the observer. Alternatively, *P_k,t,u,o_(g,d)* may take a histogram-like shape, i.e. a piecewise staircase function, in which case the effect of group size on detection probability would act on the logit-log scale in an additive manner relative to the effect of distance. With all this notation, we can then write the probability that one group went undetected as:

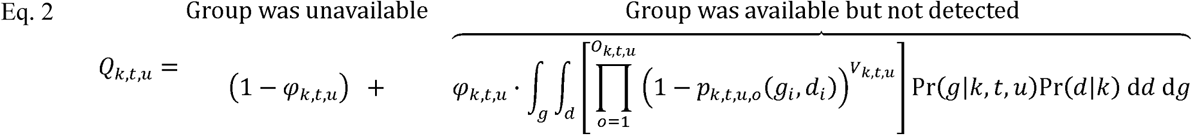

The integration over all possible group sizes and distances to the observer addresses the fact that the group size and the distance to the observer are not known but are drawn from the same distribution as the detected groups, after correcting for detection biases. In practice we computed this integral using a numerical quadrature (a.k.a. Riemann sum approximation). The probability that the total number of groups in site k during year *t* is *N_k t_* can then be expressed as a multinomial term: 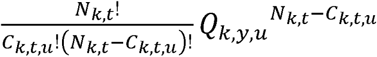, where *C_ktu_* is the number of detected groups during visit *u*.

The complete joint likelihood over all sites, years, and visits is then:

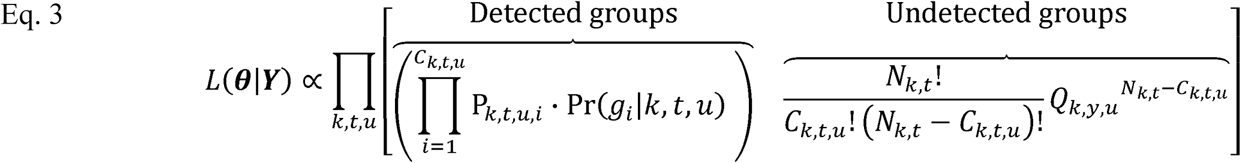

Detection and availability probabilities can be made to vary with site-specific covariates (e.g., elevation, protection status), visit-specific covariates (e.g., cloud cover, full year effect), linear temporal trends across years, or site‐ and time-random effects (the latter are however not made available in the R-package).

Eqs. 1-3 show that our model is a generalization of distance sampling. If we set all the *O_k.t.u_* and *V_k t u_* to one, if we fix all the φ_*k,t,U*_ to one, and if we remove the dependencies on *g*, we arrive at a likelihood of the form explained by Buckland, Rexstad, Marques, & Oedekoven (2015). By contrast, our likelihood cannot be simplified into a N-mixture likelihood. This is because our approach is fundamentally based on counting the groups at each site and visit, and the binomial error structure thereby applies *within*, not *across* sites and visits.

To obtain the maximum-likelihood estimates of the model parameters, we find the minimum of — log L(0|F). For that optimization we recommend the genetic algorithm with derivatives (Mebane & Sekhon, 2011), because in our experience there are many local minima in the negative log-likelihood. The preferred combination of model features should be selected using the Akaike Information Criterion (Burnham & Anderson, 2002), although to our knowledge there are no goodness-of-fit tests readily available for this type of model.

### Post-hoc manipulations

The above equations compute the number of groups *N_k t_.* To compute the population abundance, denoted *M_k,t_*, we multiplied the number of groups by the expected group size, corrected for detection biases, using the following formula:

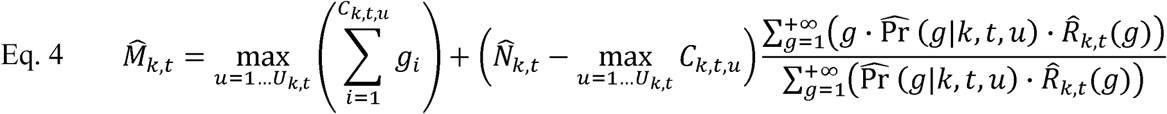

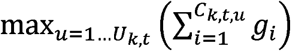 is the maximum number of individuals counted in site *k*during year *t*. *R_k_,_t_(g)* is the probability of not detecting a group of size g but of unknown distance to the observer. *R_k,t_(g)* is computed with an equation similar to Eq. 2. In practice, the sum over g was stopped after a large g chosen so that 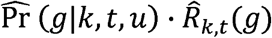 was negligible.

To estimate temporal trends in population abundance, we a posteriori regressed 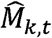 against year *t.* We considered the random effect of site *k* on the intercept, and we weighed the Poisson-distributed regression by the inverse of the sampling variance of 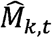. The slope of the regression represents the log-linear increase or decrease in abundance through the years.

## SIMULATION STUDIES

### Demonstrating bias in simpler methods

We considered a scenario specifically designed to challenge the IPS method but nevertheless representative of realistic sampling conditions. A population with 8 sites initially harboring 30 animals each declined by half over a 6-year period, while the expected group size increased, the probability of detection increased, and the probability of availability decreased. More precisely, the half-detection distance increased from 150 to 665m in a 1000m-long field of vision on both side of the transect, mean group size increased from 1.7 to 3.2 and had a log-scale effect of 0.5 on the half-detection distance, and availability probability decreased from 0.80 to 0.70. Combined together, these mechanisms led the expected population count to be stable over the years. We simulated 3 visits per site and per year with two observers each time. We analyzed the simulated datasets using the IPS methodology (Poisson regression of all the counts against year), using N-mixture models (one model per year, using routines in the unmarked package for R (Fiske & Chandler, 2011)), using the standard distance sampling method (one model per year and per site, using routines in the Distance package for R (Miller, 2015)), and using our new method implemented in a custom package for R (called chamois; Appendix B).

The IPS methodology failed to detect the underlying population decrease, as expected by construction of this simulation (Table 2). The N-mixture model was challenged by the small detection probability at the beginning of the simulation and the small remaining population at the end of the simulation, and as a consequence yielded many unrealistic, large population estimates, leading to unrealistic estimated rates of population change (Table 2). We note however that the number of replicates (24 per year) was in theory sufficient. The distance method was able to identify the negative population trend in most (83%) cases (Table 2), but was never powerful enough to reach statistical significance, as expected because the implementation we used did not allow the borrowing of information across years and site. The new method almost always detected the population decrease (94% of cases), but it was still not always powerful enough: statistical significance was reached in only 33% of cases (Table 2). In other words, the new method was the only one ever able to correctly detect the population decrease in the scenario we considered, but it still suffered from a high rate of type II error.

**Table 2:** Simulation study of the bias in simpler methods over 20 replicates. ‘IPS’ stands for the Poisson regression of population counts. ‘N-mixture’ means that we fit a separate N-mixture model for each year using function pcount in R-package unmarked (Fiske & Chandler, 2011). ‘Distance’ denotes the standard distance methodology: function ds in R-package Distance (Miller, 2015) applied to each year χ site combination separately. ‘Non-expected trend’ means that the estimated population trend was positive (whereas the true simulated one was negative). ‘Type I error’ means that the positive trend was statistically significant. ‘Type II error’ means that the P-value of the population trend was above 0.05, meaning that no definitive conclusion about population trend would have been reached.

|  | **New method** | **IPS** | **N-mixture** | **Distance** |
| --- | --- | --- | --- | --- |
| Non-expected trend | 17% | 83% | 50% | 20% |
| Type I error | 0% | 0% | 50% | 0% |
| Type II error | 66% | 100% | 50% | 100% |

### Finding the right bias/precision trade-off

The high rate of type II error indicates a high cost in terms of precision of the decrease in terms of bias. To investigate this potential issue, we simulated *K* sites surveyed *U* times for 6 years, by *O* observers, with 3 secondary occasions (scans). There were initially 100 animals per site. Group size and group number decreased, causing a 5% or 10% annual rate of decrease in population size (depending on scenario). Detection probability increased with group size with a slope of 0.1 on the logit-log scale but otherwise was constant over time. Since there was no temporal or spatial variation in nuisance parameters in these simulations, the IPS methodology was expected to perform best in this case. These simulations therefore quantified the loss of precision caused by increasing the number of parameters when using our new method. We ran 100 simulations per combination of K, *U* and O, and estimated the proportion of those replicates in which the population decrease was not detected.

The new method required one average more than 20 times more field effort than the IPS methodology to detect the same population trend. For example, we would need to monitor 8 sites for 6 years to be able to detect a 5% yearly rate of decrease, in a situation where only 3 sites and 3 years would be sufficient for the IPS methodology to detect the same trend (Fig. 1).

**Fig. 1:**
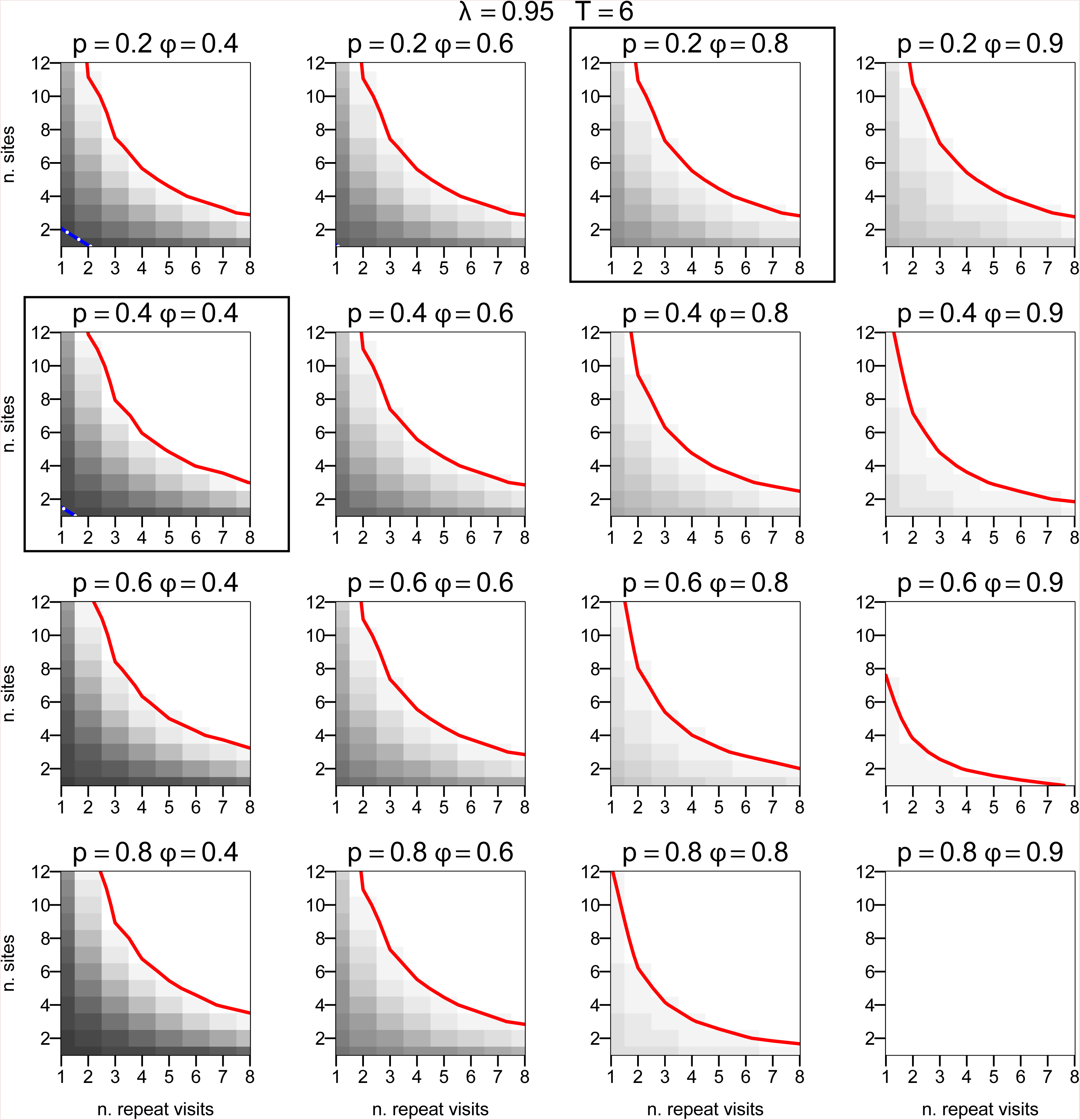
Probability of not detecting an annual rate of change (increase or decrease, at random) of 5% over 6 years, for various scenarios of detection probability *p* and availability probability *φ* using our new method. The grey shading darkens when the probability of type II error increases. The bold line is the 5% contour (right of the line, the probability of type II error is lower than 5%). The white-dashed lines correspond to the 5% contour for the population index methodology (if these white-dashed lines are absent then the probability to detect the trend was always >95% using the index). X-axis: number of repetitions. Y-axis: number of survey sites. The framed plots indicate situations that correspond to a 40% coefficient of variation, typical of mountain ungulate monitoring, even if the CV tends to get smaller than that with more replicates (Loison et al., 2006). The same figure for the probability that a 10% annual rate of change over three years was detected with a 5% risk threshold are provided in Appendix A.

These simulations also demonstrated that the new method was approximately unbiased but only when nuisance parameters took moderate values (Fig. A4c). When the availability probability was < 0.3 (meaning that more than 70% of individuals were unavailable at any time, which is extreme for an ungulate population but may be typical in e.g., marine mammals), we recorded many instabilities. The probability of availability was estimated at boundary one, i.e., over-estimated. That bias was propagated to the detection probability, which was under-estimated (Fig. A4a). This suggests that in sparse datasets only the product *φρ* might be estimable, a well-known “weak identifiability” issue that our model shares with other state-space models (Auger-Méthé et al., 2016; Barker et al., 2017). In Application case #2 (below), we demonstrate that incorporating several sources of detection data can improve the issue.

Lastly, these simulations demonstrate that the double observer protocol was never cost-effective in terms of precision compared to doubling the number of surveyed sites or the number of replicates per site.

## REAL STUDY CASES

### Application case #1: Pyrenean chamois

This case study aimed at empirically comparing the new method and the population reconstruction method. The latter is expected to perform best so is used as a reference point; the objective is to demonstrate the good performance of the new method at a fraction of the cost of the population reconstruction method. In the Bazès study area (French foothills of the Pyrenees; 43°03’N, 0°13’W), the Pyrenean chamois *(Rupicapra pyrenaica)* population has experienced a mass mortality event in the summer of 2001 that was attributed to an intoxication with an insecticide (Gibert, Appolinaire, & ONCFS SD65, 2004). Since then, breeding success has remained low, possibly due to the long life of that chemical in the environment. The monitoring program involved up to 27 visits per year since 1998. At each visit, the distance sampling protocol was applied from the same hiking trail each time. In the meantime, chamois were captured and marked every year, and then marked individuals were resighted during the population surveys.

In our new framework, we used the Akaike Information Criterion to select the presence or absence of temporal trends in detection probability, availability probability, and group size.

We also asked whether availability probability changed during the 2001 events, as would be expected if the mass mortality event was associated to a change in movement rates.

When analyzing the capture-recapture data, we used two methods. We used the Arnason-Schwarz-Gerard model (Arnason, Schwarz, & Gerrard, 1991; Schwarz & Seber, 1999) to estimate population size each year based on the year-specific estimated detection probability for marked individuals and the number of detected individuals (marked and unmarked). We also reconstructed the population trajectory using a matrix population model (Caswell, 2001) with 10 age-classes. The demographic parameters (a.k.a. vital rates) in the matrix population model were estimated from the capture-recapture data with E-Surge (Choquet, Rouan, & Pradel, 2009). Further detail can be found in Richard et al. (2017).

Availability probability was 0.57 (± standard error: 0.14) during normal years and 0.87 (± standard error: 0.74) during the 2001 intoxication event, suggesting lower movement rates. The half-detection distance varied across years between 247 and 611 meters. The lowest detection probabilities corresponded to years with staffing issues, confirming the good performance of the fit. The model without temporal variation had 14.8 AIC points more than the model with fully year-specific detection probability and an effect of the 2001 events on availability probability. Both our new method and the two capture-recapture analyses yielded the same estimated population trajectory (Fig. 2), indicating the good performance of the unmarked approach in this case relative to the much more costly mark-recapture approach. The two-way coefficients of determination (r^2^) between the year-specific population size estimates from the three methods were both 0.66.

**Fig. 2:**
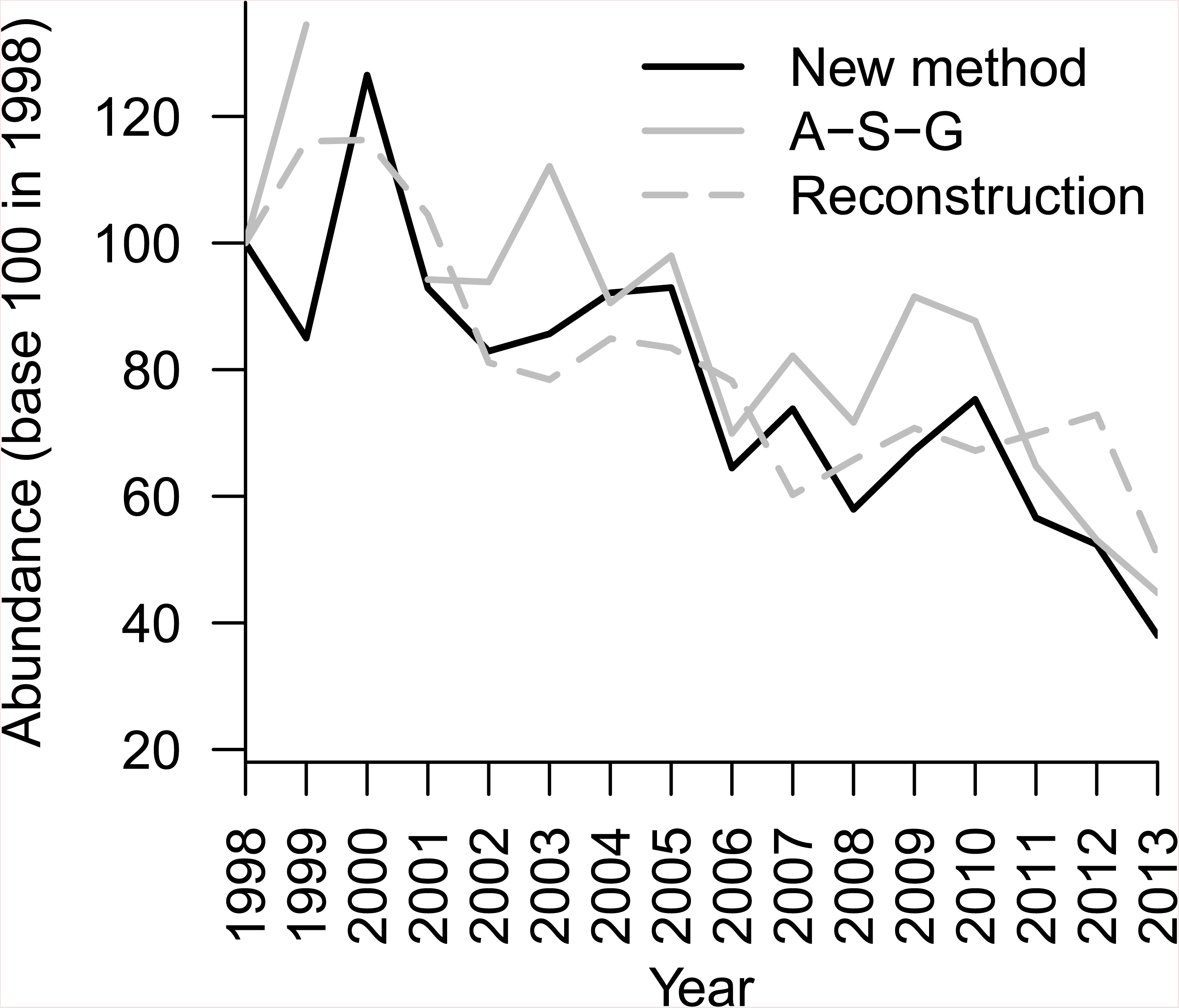
Chamois abundance in the Bazès study area estimated by the Arnson-Schwarz-Gerard model fitted to resighting data from marked animals (‘A-S-G’), by a 10 age-class population model with demographic rates estimated from individual-based data (‘reconstruction’), and by our new method.

### Application case #2: Mediterranean mouflon

We wanted to quantify how the precision of the population abundance estimate increases when we combine distance sampling, multiple-observer, and time-to-detection in a single framework, compared to when we use only one type of detection data. We also illustrated how combining multiple types of detection data can solve weak identifiability issues, namely make it possible to separate the availability probability from the detection probability.

In 2014, Mediterranean mouflons *(Ovis gmelini musimon x Ovis sp.)* were counted at three locations from fixed points in the Caroux-Espinouse national hunting and wildlife reserve (southwestern France; 43°37’54”N, 2°56’41’’E). The environment was low scrub with forest patches. On seven or eight occasions (depending on the site), two observers conducted three 15-min scans separated by 30 mins of rest. They noted which observer first recorded the animals, during which scan, and at what distance from the fixed point. We compared the standard errors of the parameter estimates when discarding the observer information, the scan information, or both.

Discarding either the time-to-detection information or the double-observer information led to a two to three-times increase in standard errors (Fig. 3). The time-to-detection information improved precision slightly more than the double-observer information. Importantly, when we discarded the time-to-detection information, the availability probability was estimated at boundary one, indicating a weak identifiability issue as described above in the simulation section. Adding the time-to-detection information solved the weak identifiability issue in this case. Based on these data we rank the observation protocols by order of increasing precision as follows: distance sampling, time-to-detection, and multiple observers.

**Fig. 3:**
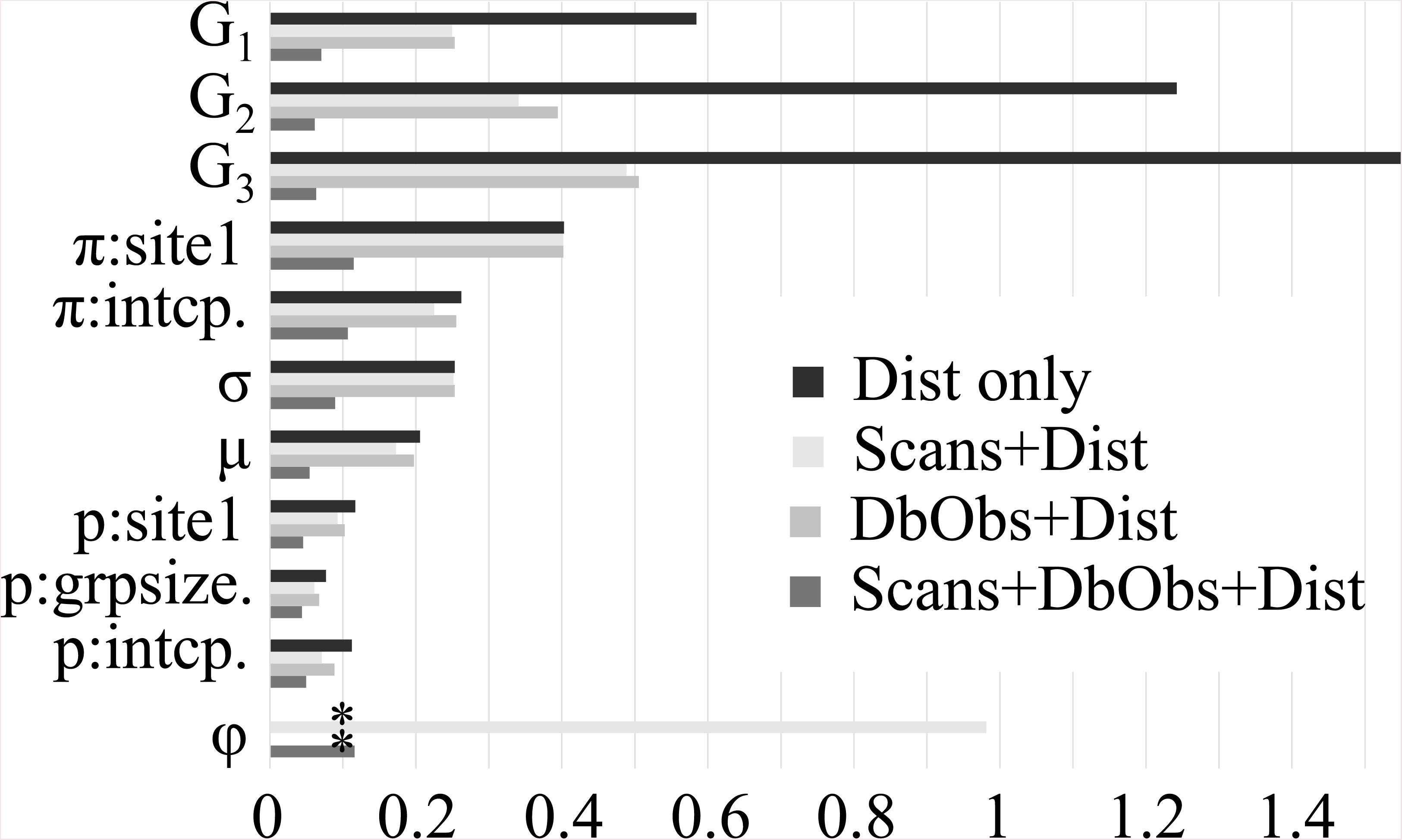
Comparison of the standard errors from our new method using various combinations of distance sampling (‘Dist’), time-to-detection (‘Scans’) and double-observer (‘DbObs’), in the mouflon case study. ‘G1’, ‘G2’, ‘G3’ stands for the log of the number of undetected animal groups in each of three survey sites, ‘π’ is the proportion of groups of size 1 (logit-scale intercept and effect of site 1). ‘σ’ is the shape parameter of the negative-binomial distribution of group sizes >1, ‘μ’ is its mean, ‘p’ is the group detection probability (logit scale intercept, effect of log-transformed group size and of site 1), and ‘φ’ is the availability probability. Asterisks indicate missing standard errors because the estimate was at boundary 1.

### Application case #3: Feral cat

This case study was chosen to illustrate the challenges associated with temporal variation in nuisance parameters and the adequate performance of the new generalized distance sampling protocol even when only distance information is available (no multiple-observer or time-to-detection). Feral cats *(Felis silvestris catus)* have been introduced to the Kerguelen archipelago (southern Indian Ocean); their abundance is a key information for a range of projects in community ecology and conservation biology. We focused on one study area (the 2.8km-long Pointe Morne transect; 49°22S,70°26E) where the cat population was surveyed on 19 occasions between 2013 and 2016 (and still ongoing) using distance sampling. At each sampling occasion, observers walked the transect back and forth until they obtained at least 30 cat sightings, later reduced to 20 sightings. They waited at least 45 minutes between the back and the forth, and at least two hours before starting again. They typically needed several days to obtain enough cat sightings. This yielded a “robust design” data structure with up to 19 secondary occasions within each of 19 primary sampling occasions. The cats could move in and out of the survey area between any two subsequent secondary occasions, but we assumed that the population abundance remained the same across secondary occasions. The population abundance could only change between primary occasions. This robust design protocol thereby departs from the terms of our model, in which we assume no exit or entries between “secondary occasions”. We thereby treated the secondary occasions as primary occasions *sensu* our model, but with a constraint of equality of the population abundance. We considered only the adult cats and did not use the information about the size of occasional family groups.

We incorporated the temporal random effect of the primary occasion into our new method, using the Gauss-Hermite quadrature within a Nelson-Mead optimization algorithm, as described in Appendix C. Random effects on the half detection distance were lognormally distributed. Random effects on availability probability were logit-normally distributed. Note that random effects are currently not available in the ‘chamois’ user interface. The proper way to select between models with and without random effects with respect to model fit and parsimony remains quite debated. We implemented a basic AIC selection procedure to select between models *(p(r)p(r), φ{·)ρ(τ)*, and φp(.)pO, where *φ* and *p* denote availability and detection probabilities, respectively, a dot denotes a time-constant model, and *r* denotes a random effect. Each random effect added a single parameter to the parameter count for the AIC.

We also applied the population index methodology (Poisson regression) and the standard distance sampling methodology (one model per primary sampling occasion). We acknowledge that the fact that primary occasions can last several days violates the assumptions of the distance sampling methodology. But our objective is actually to determine whether this represents an issue or not, by comparing the results from the standard distance sampling to the results from the new framework.

As expected, our new method yielded the same population trend as the standard distance methodology, but with better precision (Table 3: Distance vs. *φ(τ)ρ(τ)).* We obtained better precision because we borrowed information across sampling occasions and we exploited the secondary occasions, whereas in the standard distance methodology, we analyzed each sampling occasion separately without borrowing information, and we did not exploit the secondary occasions. So, applying the standard distance methodology to primary occasions that spanned over several days did not introduce a major bias, only a loss of precision compared to our new methodology. However, this loss of precision was large, meaning it could really flaw the biological inference by increasing the risk of type II error. The population index methodology underestimated the population trend compared to the other methods (Table 3: IPS vs. Distance and φ(r)p(r)). This is because of temporal variation in nuisance parameters (Table 3: φ(r)p(r) vs. *φ*(.)p(.)), which the IPS methodology did not correct for.

**Table 3:**
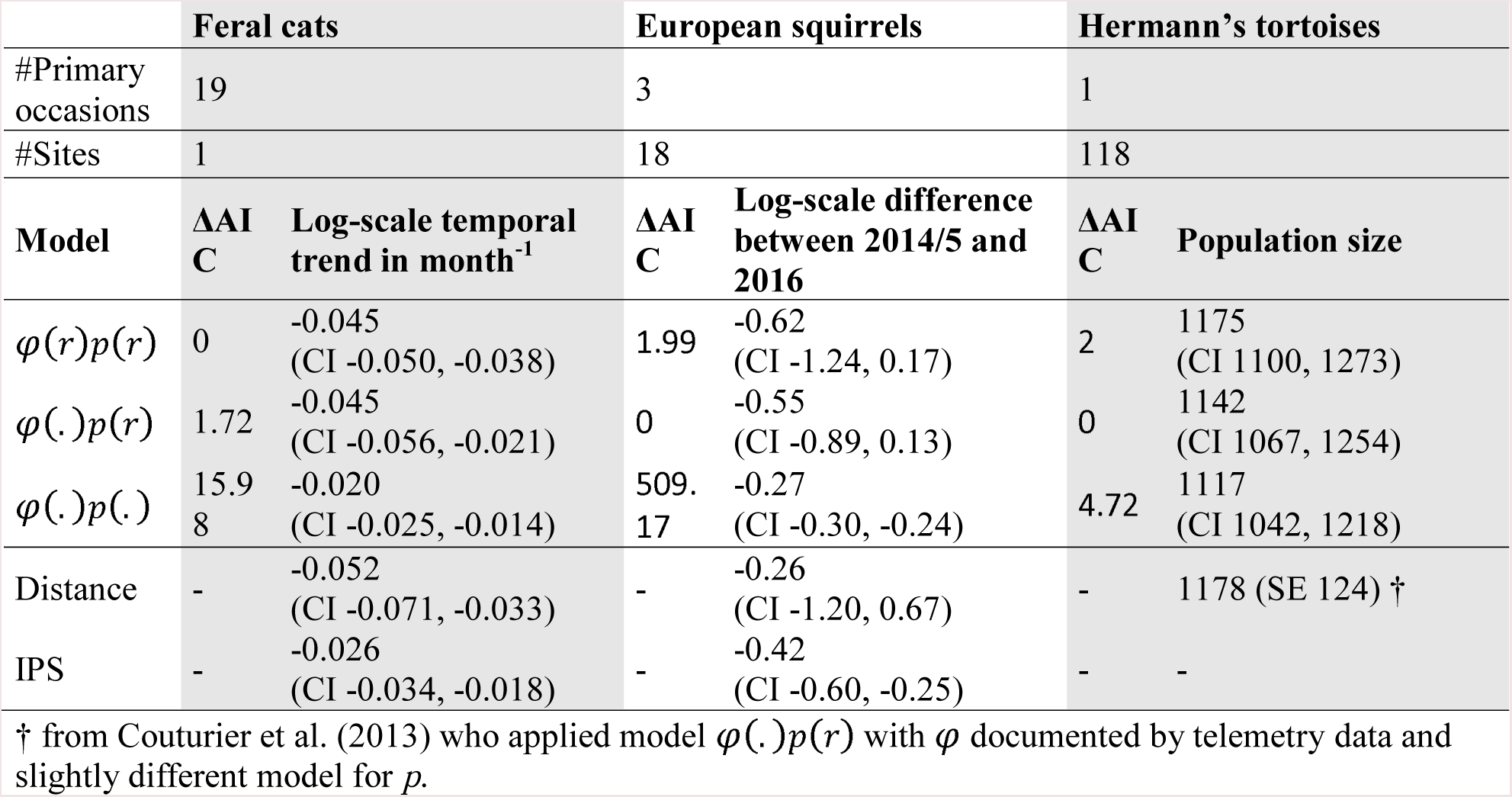
Results of the feral cat, European squirrel, and Hermann’s tortoise case studies. *φ* and *p* denote availability and detection probabilities, respectively, a dot denotes a time-constant model, *r* denotes a random effect (acting at the primary occasion scale and implemented as described in Appendix C). ‘Distance’ denotes the standard distance methodology: function ds in R-package Distance (Miller, 2015) applied to each sampling occasion separately. ‘IPS’ denotes the Poisson regression of population count against time. ÄAIC is the difference in Akaike points between the focal and preferred model. A hyphen indicates a quantity that could not be estimated. ΔAIC was not computed for Distance and IPS because the different treatment of the constant terms in the likelihoods prevented the comparison of AIC values.

|  | Feral cats |  | European squirrels |  | Hermann’s tortoises |  |
| --- | --- | --- | --- | --- | --- | --- |
| #Primary occasions | 19 |  | 3 |  | 1 |  |
| #Sites | 1 |  | 18 |  | 118 |  |
| Model | $\Delta\text{AIC}$ | Log-scale temporal trend in month<sup>-1</sup> | $\Delta\text{AIC}$ | Log-scale difference between 2014/5 and 2016 | $\Delta\text{AIC}$ | Population size |
| $\varphi(r)p(r)$ | 0 | -0.045<br>(CI -0.050, -0.038) | 1.99 | -0.62<br>(CI -1.24, 0.17) | 2 | 1175<br>(CI 1100, 1273) |
| $\varphi(.)p(r)$ | 1.72 | -0.045<br>(CI -0.056, -0.021) | 0 | -0.55<br>(CI -0.89, 0.13) | 0 | 1142<br>(CI 1067, 1254) |
| $\varphi(.)p(.)$ | 15.9<br>8 | -0.020<br>(CI -0.025, -0.014) | 509.<br>17 | -0.27<br>(CI -0.30, -0.24) | 4.72 | 1117<br>(CI 1042, 1218) |
| Distance | - | -0.052<br>(CI -0.071, -0.033) | - | -0.26<br>(CI -1.20, 0.67) | - | 1178 (SE 124) † |
| IPS | - | -0.026<br>(CI -0.034, -0.018) | - | -0.42<br>(CI -0.60, -0.25) | - | - |

† from Couturier et al. (2013) who applied model $\varphi(.)p(r)$ with $\varphi$ documented by telemetry data and slightly different model for $p$ .

### Application case #4: European squirrel

This case study illustrated how, when analyzing short time series, one needs to carefully communicate both the risk of flawed inference due to variation in nuisance parameters, and the loss of precision when incorporating more parameters in order to deal with variation in nuisance parameters. European squirrels *(Sciurus vulgaris)* were monitored for three years (2014-2016, ongoing) in the Tête d’Or urban parc (Lyon, France). Each of these 3 years, 18 transects were surveyed 6 times using distance sampling. We processed these data as the cat data, with the same three models and information-theoretic model selection, but with random effects operating at the site * primary occasion level, vs. the primary occasion level in the cat study. Squirrels results are presented along with those of the next case study below because of similarities in conclusions.

### Application case #5: Hermann’s tortoise

This case study was chosen to illustrate the effect of temporal variation in nuisance parameters on point estimates of abundance (as opposed to trend in relative abundance in the previous case studies). Hermann’s tortoise *(Testudo hermannii)* is a nationally endangered species in France with just a handful of mainland strongholds. Information about population size is important to manage the different population units and assess the effect of illegal collection for the pet trade. However, the tortoises are difficult to monitor. Records are sparse on some days and abundant on others, possibly because activity levels depend on the weather in a complex and lagged way (Couturier et al., 2013). This was expected to induce significant temporal variation in the availability probability. We compared the point estimate of abundance from our new approach to that from distance sampling with a *go* parameter estimated from telemetry, and to the estimate from N-mixture. We applied the same methodology as the cat study, with the same three models and information-theoretic model selection, but with random effects operating at the site level (vs. year or site * year level previously).

In both the squirrel and tortoise data, all the methods yielded the same conclusions, but the precision of the estimates varied significantly from one method to the next (Table 3). Adding spatial and/or temporal variation in detection probability improved the model fit and the precision of the estimates compared to the standard distance methodology (Table 3). This could have applied consequences, e.g., if managers use the lower bound of the confidence interval to inform management decisions, as is sometimes the case. However, availability probability was estimated at boundary one in both the squirrel and tortoise cases, which, following our simulation study and the mouflon case study, indicated that the probability of availability was weakly identifiable. To be able to separate the availability probability from the detection probability using survey data alone, we recommend adding a time-to-detection and multiple-observer protocol to the distance sampling protocol.

## DISCUSSION

The methods in this study build on previous efforts to jointly analyze several sources of information about the detection process of unmarked animals (Chandler et al., 2011; Fiske & Chandler, 2011; Conn et al., 2012). Our initial motivation was the analysis of mountain ungulate population surveys. This animals are gregarious and exhibit a covariation between group size, group detectability by the observer, and overall population abundance (Toïgo et al., 1996; Pépin & Gerard, 2008), which called for a model that accounted for group size effects. In addition, mountain ungulate population surveys are made extra noisy by the ease at which animals may escape detection by moving beyond the field of view of ground-based observers. This called for a model that took full advantage of all the data available (multiple observers and time-to-detection), and exploited all the patterns in between-visits variation. However, despite this initial very focused motivation, we believe the resulting framework can be relevant for a range of other surveys of group-living animals.

We proposed a fully expanded version of the model likelihood, allowing the incorporation of partially observed individual covariates. Previous “unmarked” approaches use summary statistics in a closed-form likelihood (Fiske & Chandler, 2011), which prevents the incorporation of individual and random effects. We acknowledge that this approach is computationally intensive: at least 10,000 times slower than a closed-form likelihood. It also required careful attention to local minima issues when optimizing the likelihood. Another source of concern is that the bias/precision trade-off was not always in favor of our method. However, our simulation studies clearly showed that there are situations in which our method is the only one to yield unbiased results about population trend, (Table 2). Application case #2 (mouflon) also clearly demonstrated that our new method solved a weak identifiability issue, namely made it possible to separate the availability and detection probabilities which otherwise would have been confounded.

Despite the above-mentioned loss of precision due to the increase in the number of parameter, the new method remained precise enough to be relevant for management, at least for longer studies (> 6 years) (Fig. 1). Yet, we recommend applying both the IPS methodology and our new method, maybe in a dashboard-like suite of indicators of population change. Discrepancies between the IPS and new method would then indicate either a bias in the IPS or a loss of precision in the new method. It is then possible to use a data simulation feature (provided in the chamois R-package) to run a power analysis and determine whether the loss of precision is indeed the culprit. But, from our simulations, we stress that losses of precisions remained tolerable for studies longer than 6 years.

In conclusion, we believe that our new methodological framework, a generalization of distance sampling drawing from extensive field experience counting mountain ungulates, meets several needs of wildlife managers. It controls for temporal and spatial variation in the sampling bias. It allows combining distance sampling, multiple-observer, and time-to-detection data in a flexible manner, leveraging the extra information that these different types of detection data carry, in particular regarding the ability to separate the availability and detection probabilities. It is designed to avoid type I error and presents a moderate rate of type II error once a few years of data have accumulated. We nevertheless recommend using both the IPS methodology and the new method alongside each other, because the IPS methodology is potentially biased but precise, whereas the new method is unbiased (at least for moderate amounts of nuisance) but potentially not precise enough.

## ACKNOWLEDGEMENTS

We thank Q. Richard for the capture-recapture analysis of the chamois data. We warmly thank the computing lab at IN2P3. This work was supported by the French National Park system. We thank all French national park staff for their collective insight into mountain ungulate monitoring, and in particular R. Bonet, M. Canut, J. Cavailhes, M. Delorme, T. Faivre, G. Farny, L. Imberdis, A. Jailloux, R. Papet, E. Sourp. For the mouflon and chamois studies we thank wildlife technicians J. Appolinaire, J. Duhayer, and C. Itty. For the cat study we thank the French Polar Institute (IPEV) for financial support (Program n°279) and all field workers (Y Chervaux, F. Egal, A. Lec’hvien and C. Brunet). A very warm thank you generally goes to everyone who contributed to the field efforts over the years.

## AUTHOR’S CONTRIBUTIONS

GP designed the study, wrote the R scripts, performed the statistical analyses and simulation studies, and wrote the manuscript with inputs from MG. MG designed the ungulate application cases and procured the data.

## APPENDICES

### Appendix A

Additional results from the simulation study

### Appendix B

R-package. This includes a user’s manual with installation instructions. Designed for Windows operating systems only.

### Appendix C

Using the Gauss-Hermite quadrature to fit random effect models

